# The (non) accuracy of mitochondrial genomes for family level phylogenetics: the case of erebid moths (Lepidoptera; Erebidae)

**DOI:** 10.1101/2021.07.14.452330

**Authors:** Hamid Reza Ghanavi, Victoria Twort, Tobias Joannes Hartman, Reza Zahiri, Niklas Wahlberg

## Abstract

The use of molecular data to study evolutionary history of different organisms, revolutionized the field of systematics. Now with the appearance of high throughput sequencing (HTS) technologies more and more genetic sequence data is available. One of the important sources of genetic data for phylogenetic analyses has been mitochondrial DNA. The limitations of mitochondrial DNA for the study of phylogenetic relationships have been thoroughly explored in the age of single locus phylogenies. Now with the appearance of genomic scale data, more and more mitochondrial genomes are available. Here we assemble 47 mitochondrial genomes using whole genome Illumina short reads of representatives of the family Erebidae (Lepidoptera), in order to evaluate the accuracy of mitochondrial genome application in resolving deep phylogenetic relationships. We find that mitogenomes are inadequate for resolving subfamily level relationships in Erebidae, but given good taxon sampling, we see its potential in resolving lower level phylogenetic relationships.

## Introduction

The ability to study the evolutionary histories of organisms has been revolutionized by the appearance and broad applicability of molecular methods. This ability to infer phylogenetic relationships based on molecular data was a major step forward in our understanding compared to traditional morphological comparative methods. Mitochondrial genomes offered the first possibility to use genomic scale data to infer phylogenetic hypotheses early in the history of molecular systematics. The newly accessible mitogenomic approach saw a rise in its use for resolving deep phylogenetic relationships, in arthropods and in other groups (Nardi 2003; Simon and Hadrys 2013; Song et al. 2016). Since such methods became popular, some researchers have questioned the limitations of mitochondrial genetic data for resolving early divergence events or deep phylogenetic relationships (Zardoya and Meyer 1996; Cameron et al. 2004; Talavera and Vila 2011). Nevertheless, many studies have applied mitochondrial genomes as a source of information to resolve phylogenetic relationships of varied evolutionary depth. Some studies focused on the relationships within a superorder (Cameron et al. 2009; Talavera and Vila 2011; Li et al. 2015), an order (Cameron et al. 2007; Fenn et al. 2008; Dowton et al. 2009; Kim et al. 2011; Timmermans et al. 2014; López-López and Vogler 2017; Yang et al. 2019), a family (Chen et al. 2014, 2020b; Yang et al. 2015; Li et al. 2018, 2020; Xu et al. 2020; Zhang et al. 2020) or shallower relationships like species level or even at population level.

The phylogenetic depth of a relationship affects the amount of phylogenetic signal coded in molecular data. In general, markers with higher mutation rates are only informative for the shallower evolutionary relationships or recent divergences. For clades splitting deeper in time, these fast-evolving markers will accumulate too many saturated sites and therefore tend to not resolve their phylogenetic relationship accurately. On the other hand, for markers having a very low mutation rate, a phylogenetic relationship can be too shallow for the marker to accumulate enough changes and have enough phylogenetic signal. Mitochondrial genomes usually contain relatively homogenous molecular markers in terms of mutation rate (Brower 1994). Overall, mitochondrial genes are thought to evolve relatively quickly, and e.g., the cytochrome *c* oxidase subunit I gene (COI) is used as a universal DNA barcode marker for animal species identification due to this (Hebert et al. 2003).

A peculiarity of the mitochondrial genome is the lack of recombination, which means that in practice mitochondrial DNA behaves as a single genetic marker with a unique evolutionary history. In addition, mitogenome is only maternally inherited, meaning that it has an effective population size one fourth of the nuclear genome. The mitochondrial genome is also notably susceptible to selective sweeps (Sperling 2003; Rubinoff et al. 2006). Other legitimate discordance between the mitochondrial and nuclear genome phylogenies can be associated with introgression following hybridization or retained ancestral polymorphism (Sperling 2003). Therefore, mitochondrial markers can be misleading in cases of hybridization and are more affected by demographic factors than nuclear markers.

The initial approaches to sequence the mitochondrial genomes used PCR to amplify long pieces of overlapping molecules, Sanger sequencing the long molecules and manually assembling the sequence data. The labour intensiveness and pricy nature of these methods, made the mitochondrial genomes out of reach for many research groups. With the appearance of High Throughput Sequencing (HTS) methods, the price per bp of sequencing data is dropping considerably. The advances in the methodologies to analyse the massive HTS data and the wide variety of easily accessible bioinformatic pipelines, simplify considerably their use for a large number of research groups all around the world. Therefore, it is currently easier and more economical to obtain a high number of mitochondrial genomes. The modern-day ease of sequencing mitochondrial genomes has caused a rise in the publication of single genomes practically without addressing any research question. Some authors have already started to respond to these poor scientific practices by publishing a larger number of mitochondrial genomes with a clear question at the phylogenetic depths which the markers are proven to have enough phylogenetic signal (e.g. Chen et al. 2020a).

Considering the characteristics of a mitochondrial genome as a molecular marker, the question of the phylogenetic depths whether this marker is useful arises. Also important is the question if this important genetic marker can reliably resolve phylogenetic relationships in groups which have experienced rapid radiations. In case of rapid radiations, during a short period of time, numerous lineages arise. Resolving phylogenetic relationships from past rapid radiation events is challenging due to the fact that the marker should be fast evolving enough to accumulate enough changes during the rapid radiation phase, but slow enough to not saturate the signal afterwards. One of the groups which present such challenging phylogenetic dilemmas is the moth family Erebidae.

Erebidae is one of most diverse families of moths and butterflies (Lepidoptera) with over 24,500 species described (van Nieukerken et al. 2011). In the most complete phylogenetic study of the group to date (Zahiri et al. 2012), many very short branches are recovered at the deeper levels which suggested a possible rapid radiation event. The relationships at the subfamily level within Erebidae are poorly resolved probably due to the lack of phylogenetic signal in the markers used.

Here we assemble new mitochondrial genomes of 47 species of Erebidae representing all the known subfamilies and major lineages based on the most recent phylogenetic hypotheses in order to capture all the deepest nodes within the family. In addition, downloaded 37 publicly available mitochondrial genomes and mined five transcriptomes for 11 protein coding genes found in the mitochondrial genomes. We compare the obtained phylogenetic hypotheses with known and supported relationships recovered in other studies to evaluate the phylogenetic range of accuracy of mitochondrial genomes as markers in a phylogenetic analysis.

## Material and Methods

### Taxon sampling

We sequenced low coverage whole genomes from DNA extracts of 47 species of Erebidae (Table 1). The DNA extracts were the same as those used by Zahiri et al. (2012). The taxon choice was made in order to recover all the deepest nodes within the known subfamilies and the major lineages in the family Erebidae. This allows us to focus mainly on the short deep branches which form the unresolved part of the tree for this family in published phylogenetic hypotheses. We also downloaded all the available Erebidae mitochondrial genomes from GenBank (37 genomes, Table 2), as well as mined the protein coding genes of the mitochondrion from five publicly available transcriptomes (Table 2). As outgroups we used a total of 17 taxa, which consisted of 10 Noctuidae, three Notodontidae, three Nolidae and one Euteliidae (Table 3).

**Table 1.**
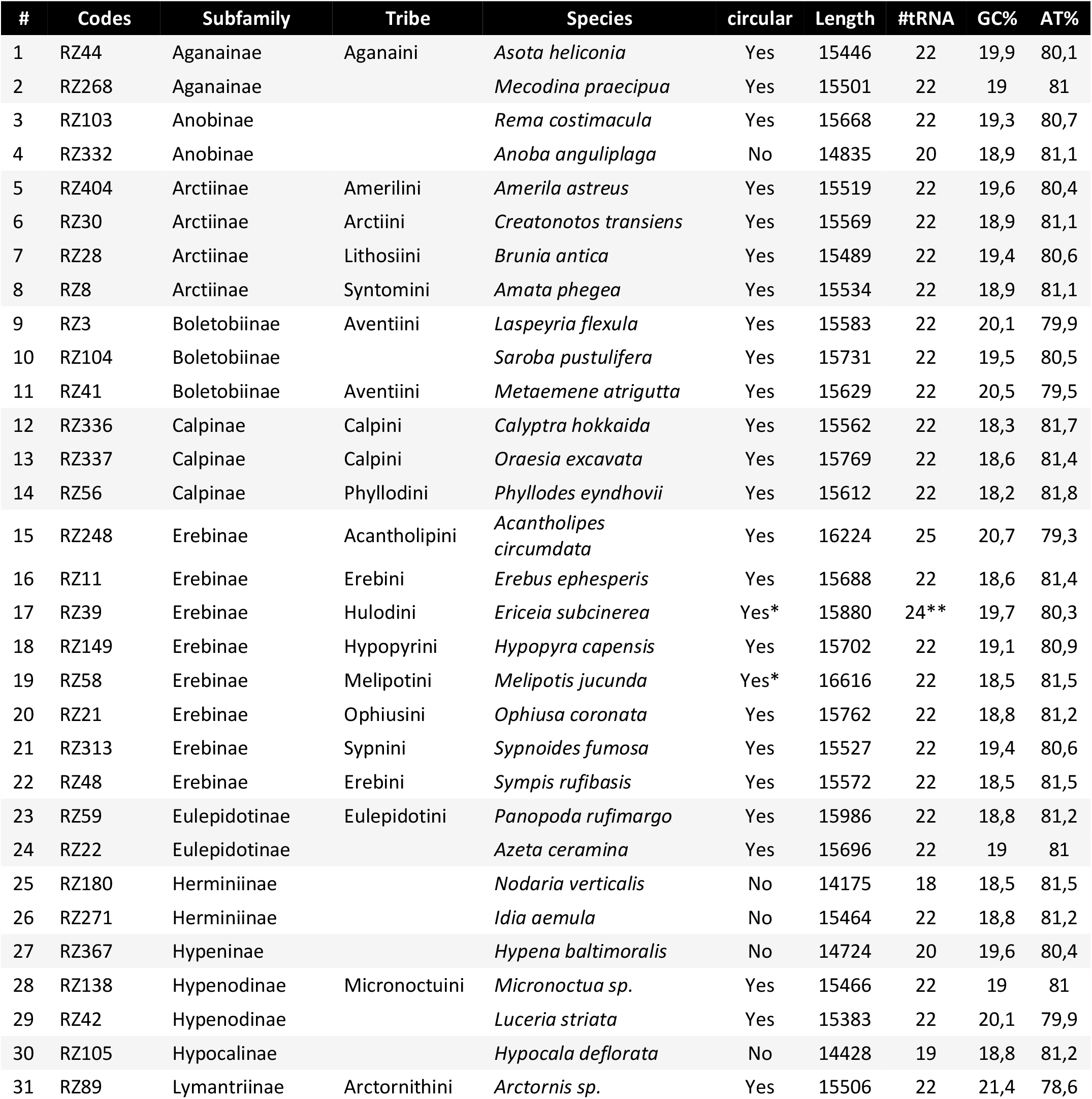
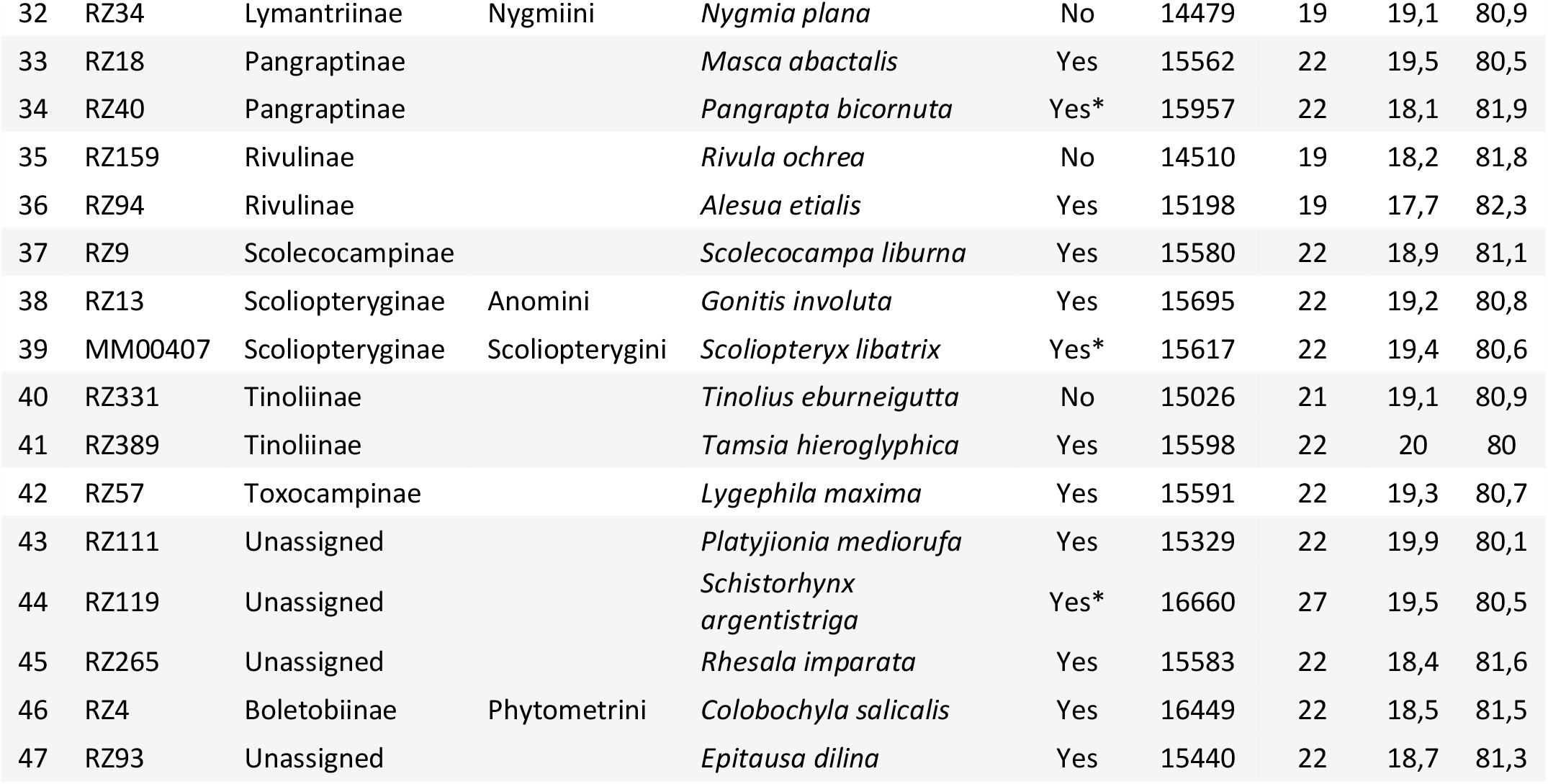
List of species sequenced in this study. The column “circular” states whether the result of Novoplasty was a circular genome (Yes) or a linear one which we manually circularized (Yes*) or not (No). Length is in base pair (bp). #tRNA is the number of tRNA recognized by MITOS. ** This genome was manually circularized, and bordering the overlapping region 2 tRNAs were repeated.

**Table 2.**
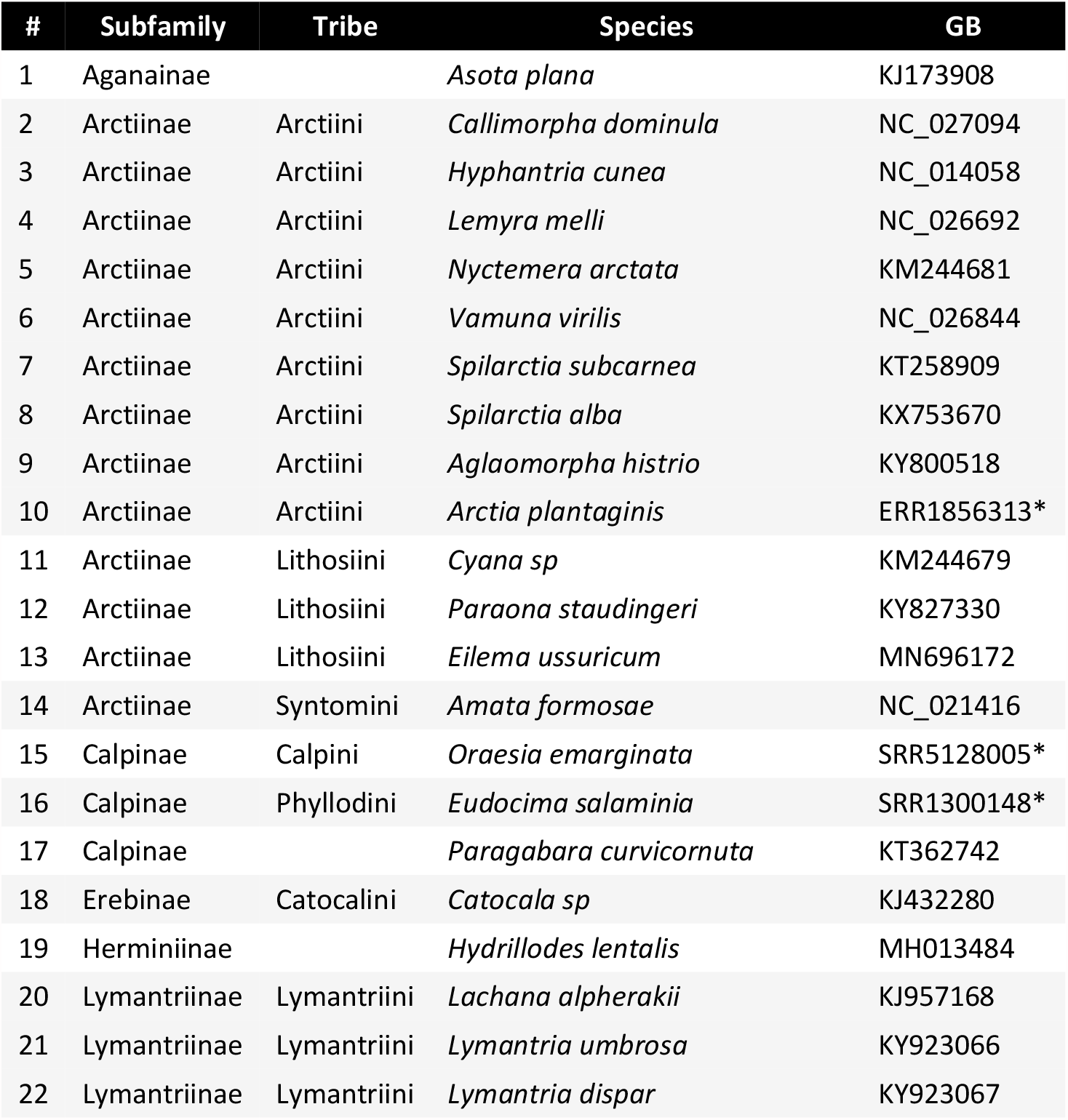
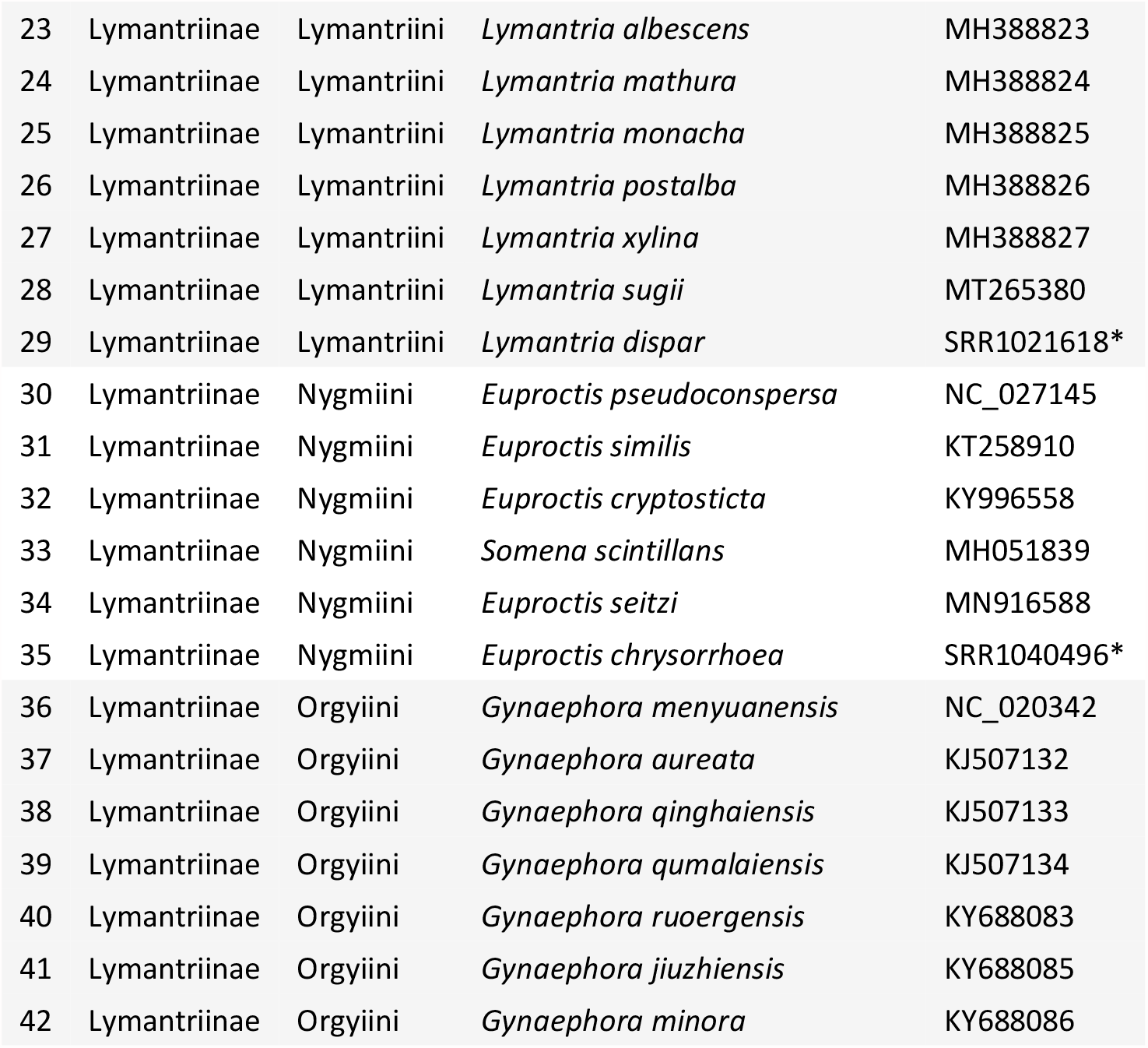
List of the Erebidae samples retrieved from other studies and their GenBank accession number (GB). Transcriptomic data used for mining of protein coding genes are marked with an asterisk (*).

**Table 3.**
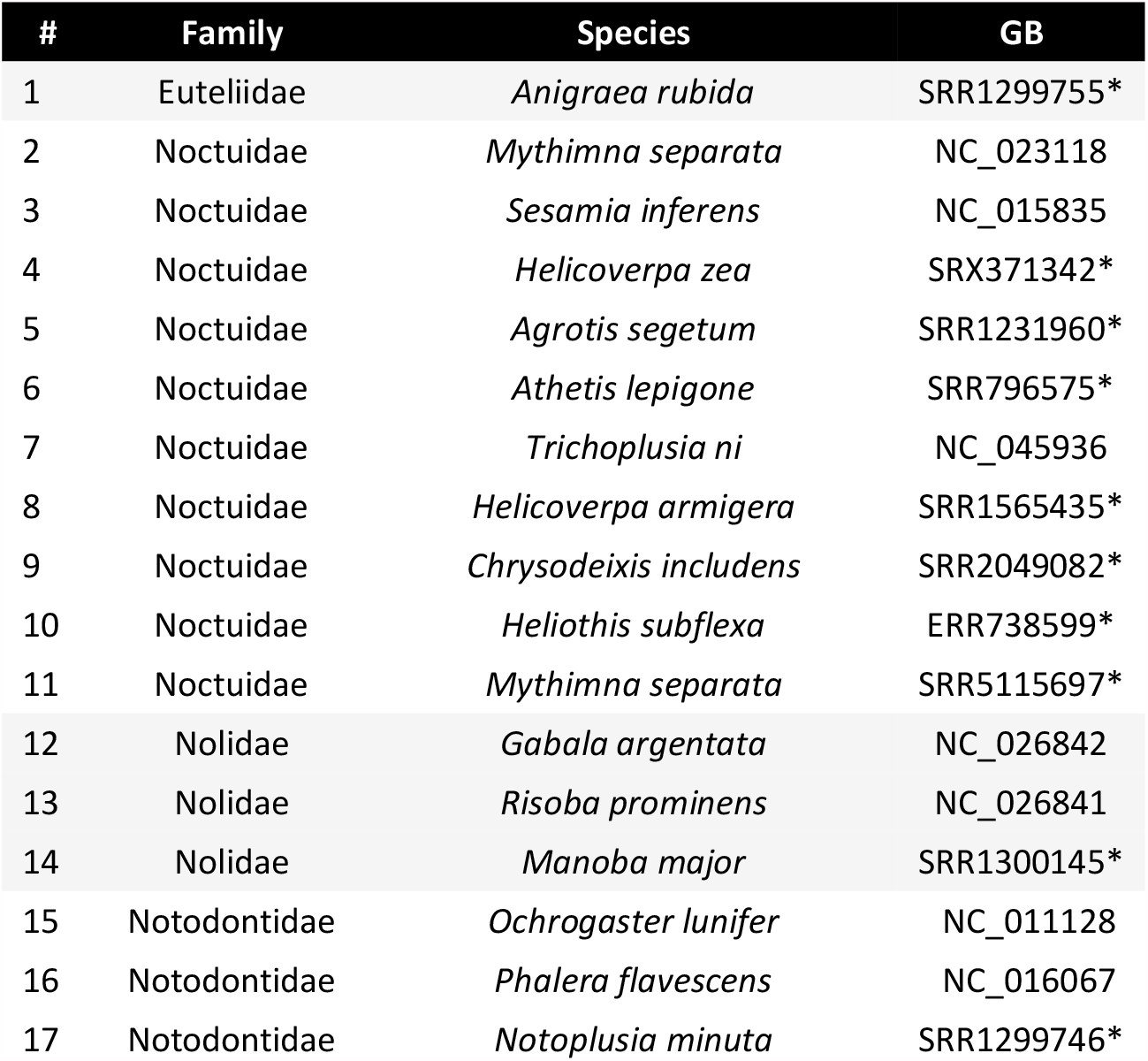
List of the Outgroups used in this study and their GenBank accession number (GB). Transcriptomic data used for mining of protein coding genes are marked with an asterisk (*).

| # | Family | Species | GB |
| --- | --- | --- | --- |
| 1 | Euteliidae | *Anigraea rubida* | SRR1299755* |
| 2 | Noctuidae | *Mythimna separata* | NC_023118 |
| 3 | Noctuidae | *Sesamia inferens* | NC_015835 |
| 4 | Noctuidae | *Helicoverpa zea* | SRX371342* |
| 5 | Noctuidae | *Agrotis segetum* | SRR1231960* |
| 6 | Noctuidae | *Athetis lepigone* | SRR796575* |
| 7 | Noctuidae | *Trichoplusia ni* | NC_045936 |
| 8 | Noctuidae | *Helicoverpa armigera* | SRR1565435* |
| 9 | Noctuidae | *Chrysodeixis includens* | SRR2049082* |
| 10 | Noctuidae | *Heliothis subflexa* | ERR738599* |
| 11 | Noctuidae | *Mythimna separata* | SRR5115697* |
| 12 | Nolidae | *Gabala argentata* | NC_026842 |
| 13 | Nolidae | *Risoba prominens* | NC_026841 |
| 14 | Nolidae | *Manoba major* | SRR1300145* |
| 15 | Notodontidae | *Ochrogaster lunifer* | NC_011128 |
| 16 | Notodontidae | *Phalera flavescens* | NC_016067 |
| 17 | Notodontidae | *Notoplusia minuta* | SRR1299746* |

### Library preparation and sequencing

In this study old DNA extracts, obtained over 10 years ago, were used to generate libraries following the protocol in Twort et al. (2020). Briefly, DNA quality was superficially checked using electrophoretic agarose gels and the high molecular weight samples were sonicated to approx. 200 – 300 bp fragments using a Bioryptor®. The DNA was then blunt-end repaired with T4 Polynucleotide Kinase (distributor), followed by a reaction clean up with the MinElute purification kit (Qiagen). This was followed by adapter ligation, reaction purification and adapter fill in. The resulting reactions were then indexed using unique dual indexes. The indexing PCR was carried out in six independent reactions to avoid amplification bias, with 15 cycles being used for each reaction. Indexing PCR reactions were pooled together prior to the final magnetic bead clean up with speedbeads. An initial bead concentration of 0.5X was used to remove long fragments that are likely to represent contamination from fresh DNA, libraries were selected with a bead concentration of 1.8X to size select the expected library range of ∼300 bp. The resulting libraries were quantified and quality checked with Quanti-iT™ PicoGreen™ dsDNA assay and with a DNA chip on a Bioanalyzer 2100, respectively.

### MtGenome assembly

In order to assemble the mitochondrial genomes (*de novo*) we have used Novoplasty (Dierckxsens et al. 2016) on the newly sequenced samples. For this analysis the raw forward and reverse read files were used with a kmer of 21. This approach gave a clean circular genome in 34 samples (72%). In an additional 5 samples (11%) the result was sufficient to manually circularize them in Geneious 10.2.6 (Kearse et al. 2012). The remaining 8 samples (17%) did not result with an assembled mitogenome using this approach probably due to their lower depth of sequencing. For these remaining samples, we used Prinseq 0.20.4 (Schmieder and Edwards 2011) to remove the reads containing ambiguous bases. We then cleaned the reads to remove low quality bases from the beginning (LEADING: 3) and end (TRAILING: 3) and reads less than 30 bp in length in Trimmomatic 0.38 (Bolger et al. 2014). Quality was measured for sliding windows of 4 bp and had to be greater than PHRED 25 on average. We then used the mirabait option in MIRA 4.0.2 (Chevreux et al. 1999, 2004) on the cleaned reads to find the reads corresponding to mitochondrial DNA. The mitochondrial reads were de novo assembled using three simultaneous approaches, the Geneious de novo assembler, SPAdes assembler 3.10.0 (Nurk et al. 2013) and plasmidSPAdes (Antipov et al. 2016), all of which are implemented in Geneious. For each sample, all contigs over 500 bp were aligned to a reference mitochondrial genome of *Lymantria dispar* (Erebidae). The consensus sequence of the alignment was then used as a reference to map the mitochondrial reads in Bowtie2 (Langmead and Salzberg 2012) as implemented in Geneious with default parameters. All the resulting assembled genomes were annotated using MITOS (Bernt et al. 2013).

### Phylogenetic analyses

Eleven protein coding genes (PCG) were extracted from all mitochondrial genomes. This dataset includes the genes coding for ATP synthase membrane subunit 6 (*ATP6*), cytochrome *c* oxidase subunit I to III (*COI-III*), cytochrome b (*Cytb*), NADH dehydrogenase 1 to 5 (*ND1* - *ND5*) and the NADH-ubiquinone oxidoreductase chain 4L (*ND4L*). We excluded two genes (*ATP8* and *ND6*) from our dataset as they did not align properly. Each gene was aligned separately using MAFFT v7.450 (Katoh 2002; Katoh and Standley 2013) as implemented in Geneious with default options. The sequences were curated and maintained in VoSeq (Peña and Malm 2012), after revision and manual correction of the alignments. Using the VoSeq database application, we created a nucleotide concatenated dataset (nc) with a total length of 10,245 bp and an amino acid dataset (aa) of 3,415 characters.

We ran maximum likelihood (ML) analyses with both nc (partitioned by gene and codon position) and aa (partitioned by gene) datasets using IQ-TREE 2.0.6 (Nguyen et al. 2015). In both analyses the best substitution model and partitioning scheme was selected by ModelFinder (Kalyaanamoorthy et al. 2017) with “-m MFP+MERGE” option. We evaluated the node supports with 5000 ultrafast bootstrap approximations (UFBoot2) and 1000 SH-like approximate likelihood ratio test (Guindon et al. 2010; Hoang et al. 2018) using the “-B 5000 - alrt 1000” option. We used the “-bnni” option to reduce the risk of overestimating branch supports in ultrafast bootstrap approximation analysis. Additionally, we tested the best partitioning scheme for the nucleotide dataset partitioned by gene only in PartitionFinder2 (Lanfear et al. 2017). In this analysis we limited the tested models with the option “models = mrbayes”. The obtained partitioning scheme was used to perform a Bayesian phylogenetic analysis in MrBayes 3.2.7 (Ronquist et al. 2012). This analysis ran for two independent runs of 10^7^ generations sampling every 10^3^ steps. This analysis was repeated five times. The convergence of the runs was checked in Tracer 1.7.1 (Rambaut et al. 2018). The resulting trees were visualized and rooted in FigTree v1.4.3 (Rambaut 2016) using the outgroups. The COI gene was extracted from all the assembled genomes to compare with the sequences obtained with Sanger sequencing as an extra quality control.

The software Mira and Novoplasty were run using the resources provided by SNIC through Uppsala Multidisciplinary Center for Advanced Computational Science (UPPMAX) under Project SNIC 2018-8-347. The software PartitionFinder2 and MrBayes were run using the CIPRES Science Gateway infrastructures (Miller et al. 2010). The raw whole genome data is deposited in GenBank under the BioProject number PRJNAXXXXX. All data in the supplementary material, the alignment, the annotated genomes and the results can be found and downloaded from the GitHub repository: github.com/Hamidhrg/ErebidMtGenome.

## Results

From the total number of 47 obtained genomes, 34 were fully assembled as circularized genomes. For the base frequency and basic genome composition result we only focus on the 34 good quality genomes. They varied in length from 15,198 bp in *Alesua etialis* (Rivulinae) to 16,449 bp in *Colobochyla salicalis* (Boletobiinae, Phytometrini). Their AT base frequency ranged between 78.6% in *Arctornis* sp. (Lymantriinae) to 82.3% in *Alesua*. Their tRNA number was between 19 in *Alesua* to 25 in *Acantholipes circumdata* (Erebinae) (Table 1). The annotated genomes are available through our online GitHub repository.

ModelFinder in IQ-Tree2 merged the 33 possible partitions of the nucleotide dataset into 13 and found their corresponding best substitution models (Table 4). The partition sizes ranged between 96 to 1,411 bp (788 bp mean partition size). In total the dataset included 4,789 phylogenetically informative sites.

**Table 4.** ModelFinder best partitioning scheme. Length column corresponds to the total number of base pairs forming the partition. “Infor” and “Invar” stand for number of informative and invariable sites respectively.

| Partition | Markers | Length (bp) | Infor | Invar | Model |
| --- | --- | --- | --- | --- | --- |
| 1 | ATP6_pos1, COII_pos1, COIII_pos1, CytB_pos1 | 1083 | 397 | 592 | GTR+F+R7 |
| 2 | ATP6_pos2, COI_pos2, COII_pos2 | 965 | 139 | 758 | GTR+F+R7 |
| 3 | ATP6_pos3 | 227 | 197 | 15 | TPM2+F+I+G4 |
| 4 | COI_pos1 | 510 | 130 | 345 | GTR+F+I+G4 |
| 5 | COI_pos3, COII_pos3, ND3_pos3 | 847 | 580 | 227 | TIM+F+R5 |
| 6 | COIII_pos2, CytB_pos2, ND2_pos2, ND3_pos2 | 1039 | 218 | 705 | TVM+F+R4 |
| 7 | COIII_pos3, CytB_pos3 | 628 | 572 | 29 | TPM3+F+R7 |
| 8 | ND1_pos1, ND4_pos1, ND4L_pos1, ND5_pos1 | 1411 | 599 | 635 | GTR+F+R5 |
| 9 | ND1_pos2, ND4_pos2, ND4L_pos2, ND5_pos2 | 1411 | 310 | 950 | GTR+F+R5 |
| 10 | ND1_pos3, ND4_pos3, ND5_pos3 | 1315 | 1090 | 93 | TIM+F+R7 |
| 11 | ND2_pos1, ND3_pos1 | 411 | 210 | 133 | GTR+F+R5 |
| 12 | ND2_pos3 | 302 | 269 | 13 | GTR+F+I+G4 |
| 13 | ND4L_pos3 | 96 | 78 | 4 | GTR+F+I+G4 |
| **Total** |  | **10245** | **4789** | **4499** |  |

The ML analysis of the nc dataset resulted in the best resolved tree (Figure 1). The family Erebidae was a well-supported monophyletic group. All the other families used as outgroups were also recovered as monophyletic with more or less high support. Within Erebidae, most of the subfamilies that had more than one representative were found to be monophyletic, including Lymantriinae, Arctiinae, and Erebinae. A few subfamilies did not form monophyletic groups, such as Pangraptinae and Aganainae.

**Figure 1.**
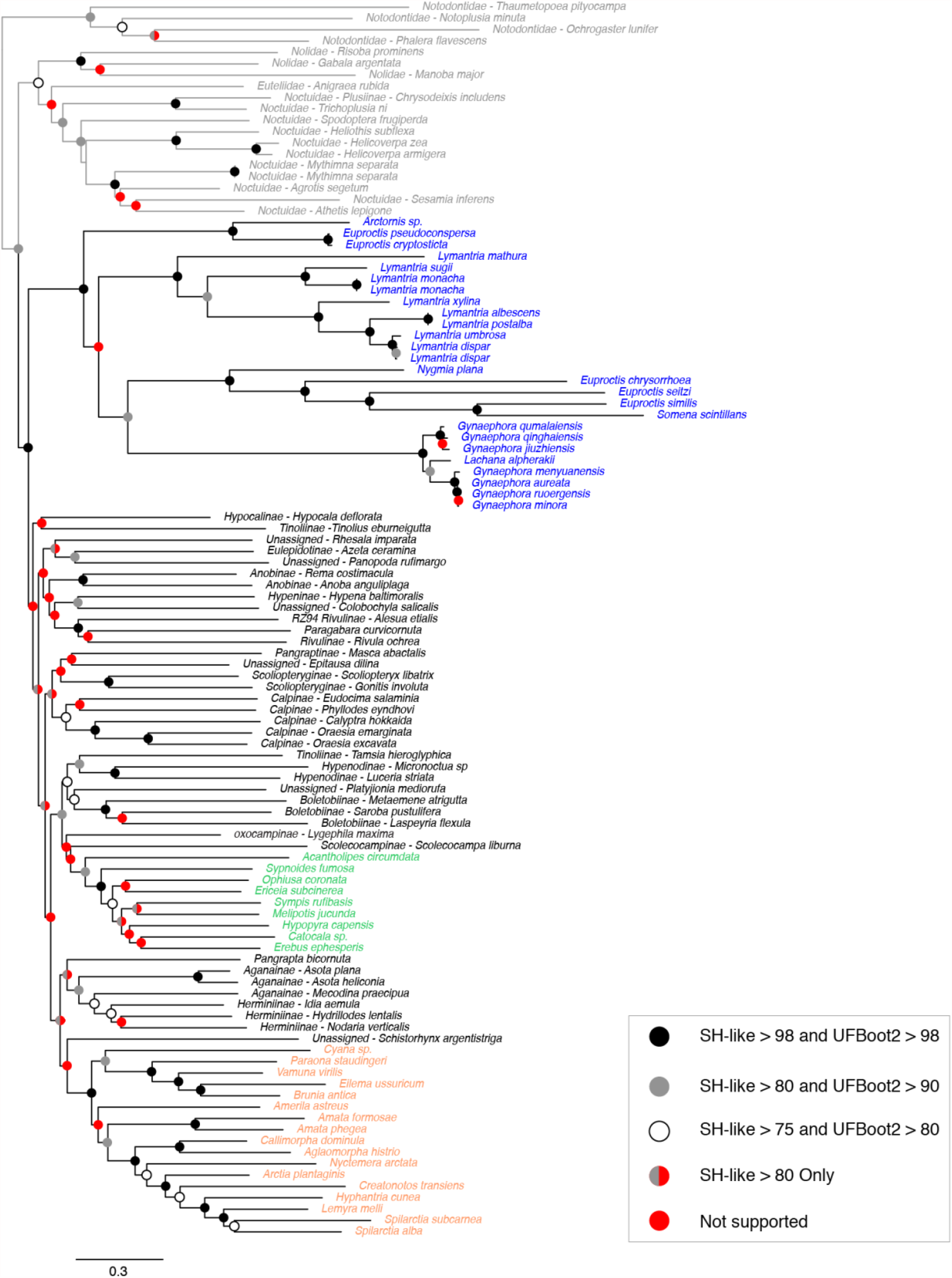
The ML tree obtained using the nc dataset in IQ-Tree2. The species names coloured in blue corresponds to the subfamily Lymantriinae, green Erebinae and orange Arctiinae. Black circles represent highly supported nodes, grey supported nodes, white low support and red not supported nodes. The outgroup lineages are coloured in grey.

In contrast the ML analysis of the aa dataset resulted in very anomalous trees (supplementary material). First of all, it appeared very sensitive to missing data. Therefore, 3 samples with the highest amount of missing data were deleted from the dataset and a new analysis was run. The resulting tree improved slightly, however it was still very anomalous. In the case of the Bayesian inference (BI) in MrBayes using the nc data partitioned according to the PartitionFinder2 analysis, all of the ten chains (five runs of 2 independent chains) reached the stationary phase but none of the runs converged with each other. The analysis was repeated for a longer (up to 10^8^) generation number and with a higher temperature (up to temp = 0.7) resulting in the same issue.

## Discussion

The most comprehensive study focused on the phylogenetic relationships of Erebidae up to date was published by Zahiri et al. (2012). Using seven nuclear and one mitochondrial markers (for a total of 6,407 bp) they inferred a phylogenetic hypothesis with numerous unsupported short branches which did not resolve the relationship among different subfamilies and tribes. Similarly, we find that mitochondrial genomic data were not able to resolve the relationships of subfamilies with any confidence. We do find that the family itself is a strongly supported monophyletic group, with respect to the outgroups. As earlier studies have found (Zahiri et al. 2011; Regier et al. 2017), our results also showed that the relationships among the other four lineages (Notodontidae, Nolidae, Euteliidae, and Noctuidae) are not clear, although they do form a monophyletic assemblage with good support. Within the quadrifid noctuoids (Noctuidae, Nolidae, Euteliidae, and Erebidae), the sister group of Erebidae remains unresolved. Our phylogeny placed Erebidae as the sister to the other quadrifids (Nolidae, Euteliidae, and Noctuidae), however, (Zahiri et al. 2012) found Euteliidae+Noctuidae in this position in their ML analysis and Noctuidae as sister to Erebidae in parsimony.

We find that the subfamily Lymantriinae is sister to the rest of Erebidae, however this position has no support. Zahiri et al. (2012) did not find Lymantriinae in the same position but also in that study its position is not supported. Fibiger & Lafontaine (2005) placed Lymantriinae adjacent to Arctiinae to reflect the close association found by Mitchell et al. (1997, 2000). They also noted that arctiines and lymantriines are not basal clades but appear to be highly specialized lineages derived from within Erebidae. Lymantriinae like Herminiinae, Aganainae and Arctiinae shared a unique apomorphic character — a prespiracular counter-tympanal hood — that had been interpreted as a plesiomorphic condition in quadrifid Noctuoidea for a long time. Our results did not support such a relationship. Branch lengths within the Lymantriinae clade appear to be longer than in the rest of the tree (Fig. 1). This pattern of exceptionally long branch lengths among Tussock moths were also observed by (Zahiri et al. 2012). One reason for this would be a higher rate of molecular evolution within Lymantriinae, although the reasons for a higher rate are not known at the moment. The support values of the nodes in this clade could appear high at first, but after further attention it is clear that the high support values only correspond to the relationships within the same genus and not between the different genera. Wang et al. (2015) studied this subfamily using eight molecular markers, and they found that the relationships between different tribes are poorly supported. We did not include the tribe Daplasini which (Wang et al. 2015) found to be sister to the rest of the subfamily. We found the tribe Arctornithini to be sister to the rest of the Lymantriinae species that we had sequences for, a result which is in concordance with (Wang et al. 2015).

Our phylogeny placed the representative of Pangraptinae (*Pangrapta bicornuta*) as sister to Aganainae and Herminiinae (Fig. 1). This triplet (Pangraptinae and Aganainae + Herminiinae) is weakly associated with a clade (with no support) consisting of the enigmatic genus *Schistorhynx* Hampson and Arctiinae. The study of Zahiri et al. (2012) placed Pangraptinae within the group of subfamilies with prespiracular counter-tympanal hoods, although morphological examinations of various genera of pangraptine revealed that they have a typical erebine postspiracular hood. Zahiri et al. (2012) concluded that the prespiracular feature can be either the result of convergent evolution in Lymantriinae and the clade comprising Herminiinae, Aganainae and Arctiinae, or as a unique derivation in the larger clade that encompasses all these groups, with subsequent reversal in Pangraptinae. Our results do not support any of these hypotheses since Lymantriinae, Pangraptinae, Herminiinae, Aganainae and Arctiinae are not recovered as a monophyletic group. Instead, our results suggest independent evolution of prespiracular counter-tympanal hoods in Lymantriinae and a clade of four subfamilies Pangraptinae, Herminiinae, Aganainae, Arctiinae with subsequent reversal in Pangraptinae.

The relationships within Arctiinae are better supported and appear to be better resolved. The clade composed by *Cyana* sp., *Paraona staudingeri, Vamuna virilis, Eilema ussuricum* and *Brunia antica*, representing the tribe Lithosiini, is placed as the sister group to the rest of the subfamily. This position of Lithosiini is in concordance with previously published studies (Zahiri et al. 2012; Zaspel et al. 2014; Rönkä et al. 2016; Dowdy et al. 2020). Also, the position of *Amerila astreus* (Amerilini), even though it is not supported, and the relationship of Callimorphina and Arctiina subtribes are similar to the afore mentioned studies.

Erebinae is recovered as a monophyletic group, however, within the subfamily there is a lack of support for the resolution of the relationships among different genera. The placement of *Acantholipes circumdata* (Acantholipini) as the sister group of the rest of the subfamily is also recovered in (Zahiri et al. 2012) and Homziak et al. (2019). Homziak et al. (2019) used anchored hybrid enrichment (AHE) phylogenomics to resolve the deep node relationships within this subfamily. The position of the species *Sypnoides fumosa* (Sypnini) in Erebinae clade is in concordance with Zahiri et al. (2012) and Homziak et al. (2019). The rest of the relationships within the subfamily are poorly resolved and do not agree with the mentioned studies.

Boletobiinae, Calpinae, Scoliopteryginae, Rivulinae, Hypenodinae, and Anobinae are recovered as monophyletic entities, however, their interrelationships are either weakly supported or have no support in our analysis. The relationships among the subfamilies have not been resolved in any published phylogenetic work up to date. Zahiri et al. (2012) suggested that the short internal branches connecting different subfamilies and some tribes are potentially due to a rapid radiation. Therefore, more data and more comprehensive taxon sampling are needed in order to resolve these relationships. The results of our study show very low support values for these internal nodes suggesting that the amount of information coded in the mitochondrial genome is not sufficient to deal with such rapid radiations of similar or older ages. One of the caveats of our study is the sporadic taxon sampling in our dataset. Although our dataset has low taxon sampling, it is still comparable to most multi-locus phylogenetic studies in number of species and definitely larger than most phylogenomic datasets. Hence, we believe that expanding the taxon sampling will definitely improve the phylogenetic resolution. Nevertheless, most probably, it will only affect the more recent divergence events at genera and species level as is visible in the better sampled clades in our study (e.g., Arctiinae and Lymantriinae).

In the dataset studied here, the amount of phylogenetic signal coded in the mitochondrial genome was not sufficient to resolve satisfactorily the relationships among the representatives of the Erebidae family. This is especially visible in the deeper nodes (Figure 1). The lack of resolution for deep and short branches suggest that on one hand, this dataset has probably relatively high mutation rates which cause saturation issues for deepest relationships, and on the other hand, the amount of coded signal in mitochondrial genome, seems to not be adequate to recover deep short branches. Meaning that while higher mutation rates are theoretically beneficial to recover short speciation events in general, the small size of this marker does not allow to store enough signal to counter the effect of saturation of the signal in time.

## Conclusion

The advances in sequencing technologies and the bioinformatics supporting it have revolutionized molecular systematics, evolutionary biology and phylogenomics, among other fields. Especially with the advances in HTS, sequencing a large number of mitochondrial genomes is relatively cheap and does not need much more infrastructure than the traditional PCR lab. This has allowed a rise in the number of new mitochondrial genomes being published practically on a weekly base in the last few years. These short publications usually publish a single new mitochondrial genome together with a very brief and rudimentary phylogenetic analysis.

In this study we question the utility of mitochondrial genome data to resolve deeper phylogenetic relationships accurately, or to resolve relationships of groups involving rapid radiation events. Based on our findings, at least for the erebid moths, mitochondrial genomes are not a good enough source of information to resolve the relationships within and between subfamilies. The relationships between different close tribes could potentially be studied with a high enough taxon sampling in Erebidae. We also show that it is clear that amino acid datasets based on mitochondrial protein coding genes are not useful to study phylogenetic relationships at this level.

## Acknowledgements

HG received funding from the European Union’s Horizon 2020 research and innovation program under the Marie Skldowska-Curie grant agreement No. 6422141. NW acknowledges funding from the Swedish Research Council (Grant No. 2015-04441). We thank Marko Mutanen for sending us the DNA extract for *Scoliopteryx libatrix*.

